# Automated Labeling of Scientific Names and Etymological Trend Analysis in Phytophagous Arthropods Using Large Language Model

**DOI:** 10.1101/2025.03.16.643572

**Authors:** Kota Nojiri, Keito Inoshita, Haruto Sugeno

## Abstract

Scientific names, especially epithets, are derived from various factors, not only species characteristics but also cultural backgrounds, such as the names of people. They reflect how species were perceived at the time. However, several ethical issues have been raised, such as naming species after criminals and gender imbalance in eponyms (epithets named after people). Previous research has been conducted through thorough literature reviews with random sampling, which requires significant time and effort. In this study, the accuracy of the automated labeling using a Large Language Model (LLM) was assessed, and the temporal etymological trends of 2,705 species of phytophagous arthropods were investigated. LLM-based classification achieved F1 scores above 75% and accuracy above 90% in the *Morphology, Host, Geography*, and *People*. However, the *Ecology & Behavior* and *Other* exhibited accuracy issues. Analyses using the Generalized Additive Model (GAM) revealed shifting naming trends, with a decrease in *Morphology* and an increase in *Geography* and *People*, consistent with previous research on spiders. This study demonstrates the effectiveness of LLM-based classification for epithets and provides a new perspective on the social and scientific debates surrounding scientific names based on etymological trends.

## INTRODUCTION

Humankind needs to name natural phenomena and objects to understand them. Systematic naming theories have been established in various ways (e.g., the periodic tables of the elements, and the nomenclatures of stars), forming the foundation of science.

In the naming of animals, the binomial nomenclature proposed by Carl Linnaeus about three centuries ago remains widely used in modern biology. It consists of a generic name and a specific name/epithet, representing taxonomic and species characteristics. The International Code of Zoological Nomenclature (ICZN) is widely accepted as the standard for naming animals. The authors have the freedom to name newly discovered species, provided that they adhere to minimum rules set by the ICZN.

The origin of species epithets is primarily derived from morphology, ecology, distribution, eponyms, and cultural background(Figueiredo and Smith, 2010; Jasper et al., 2021; Macêdo et al., 2023; Mammola et al., 2023). In parasitic animals or species heavily dependent on other organisms, epithets are sometimes named after their hosts(Poulin et al., 2022). While these epithets are valuable for scientific purposes, several issues regarding naming have been raised. For instance, ethical concerns have been pointed out regarding eponyms honoring racists or criminals(Guedes et al., 2023; Roksandic et al., 2023), and gender imbalance, with about 90% of eponyms commemorating men(Figueiredo and Smith, 2010; Poulin et al., 2022; Vendetti, 2022). Some taxonomists argue that these problems stem from the ICZN’s principle of priority (where the oldest valid name must be applied to a scientific name) and the historical use of Latin (though now largely obsolete)(Gillman and Wright, 2020; Roksandic et al., 2023; Rummy and Rummy, 2021; Wright and Gillman, 2022). However, revising the ICZN would require renaming certain species, potentially leading to inconsistencies in scientific classification. Thus, opposing views have also been expressed(Antonelli et al., 2023; Ceríaco et al., 2023; Jiménez-Mejías et al., 2024; Orr et al., 2023; Pethiyagoda, 2023; Thiele, 2023). Moreover, scientific names influence which species are selected as research subjects (Mlynarek et al., 2023). Above all, the significant impact of scientific names on both society and science is widely recognized, and related research has been actively conducted in recent years (Figueiredo and Smith, 2010; Jasper et al., 2021; Macêdo et al., 2023; Mammola et al., 2023; Mlynarek et al., 2023; Poulin et al., 2022; Vendetti, 2022).

Previous etymological research has been conducted through thorough literature reviews and cross-validation using random sampling(Figueiredo and Smith, 2010; Jasper et al., 2021; Macêdo et al., 2023; Mammola et al., 2023; Mlynarek et al., 2023; Poulin et al., 2022). However, this method requires substantial time and effort. Given that it is estimated that over 80% of arthropods remain undiscovered (Stork, 2018), it is not practical to invest excessive resources in analyzing the scientific names of already identified species. Priority should be given to describing new species and conserving the environment, making it necessary to develop effective and efficient methods for etymological analysis.

Recently, the Large Language Model (LLM) has achieved human-level performance in text classification(OpenAI et al., 2024). LLM can extract the semantic meaning from text by learning from vast amounts of textual data(Wei et al., 2022). It has been suggested that automated labeling using LLM could significantly reduce the time and cost of classifying the origins of the epithets(Inoshita et al., 2025). However, previous research has been limited to spider taxa.

This study was conducted with the following objectives: 1) To evaluate the accuracy of LLM-based labeling in arthropods and demonstrate that LLM classification can be applied beyond spiders. 2) To elucidate the temporal etymological trends of phytophagous arthropods and compare them to other taxa.

## MATERIALS AND METHODS

We used an open-source etymological dataset of phytophagous arthropods (Heard et al., 2023). It comprises 30 species that are frequently selected as research subjects for studying Host-Associated Differentiation (HAD), along with their respective genera, totaling 2,709 species. Each species was originally classified into seven categories: *Morphology, Host, Place, Behavior, Habitat, Person*, and *Other*. To align with previous research (Inoshita et al., 2025), we reclassified them into six categories: *Morphology, Host, Geography, Ecology & Behavior, People*, and *Other*. Specifically, we renamed *Place* as *Geography, Person* as *People*, and merged *Behavior* and *Habitat* into *Ecology & Behavior. Morphology* refers to names based on morphological characteristics such as size, shape, and color. *Host* refers to names related to their host taxa. *Geography* refers to names based on species distribution or the site where they were found. *Ecology & Behavior* refers to names based on their habitat and behavioral traits. *People* refers to names honoring individuals, such as scientists. *Other* includes names that do not fit into the above categories. In the dataset, four species lacked published years, so they were omitted from this study, resulting in a final analysis of 2,705 species. For labeling, GPT-4o mini was selected as the LLM, accessed through OpenAI’s API. We applied the prompt provided by Inoshita et al. (2025), modifying it with a few-shot example to match our target.

R version 4.4.1(R Core Team, 2024) was used for data processing and analysis throughout the experiment. To evaluate the similarity of the Human-based and LLM-based classification, we compared the total count for each year and calculated the Pearson Correlation Coefficient (PCC) and Median Absolute Error (MAE). Subsequently, the Generalized Additive Model (GAM) was applied to analyze the temporal trends in etymology. Additionally, a GAM integrating Human-based and LLM-based labeling was constructed to compare time-series changes across both methods. All GAM models assumed a quasibinomial distribution and logistic regression was employed.

## RESULTS

Table 1. shows the performance of LLM-based classification. The highest accuracies were found in *Ecology & Behavior* (94.5%) and *Geography* (94.1%). *People* (93.0%) and *Host* (90.3%) also achieved over 90%, while *Morphology* (88.3%) and *Other* (85.0%) exhibited high accuracy as well. F1 scores were high in *Morphology* (82.6%), *People* (80.0%), *Host* (76.4%), and *Geography* (75.8%), indicating an appropriate balance between precision and recall. On the other hand, *Other* (38.3%) and *Ecology &* B*ehavior* (31.7%) had low F1 scores.

**Table 1.**
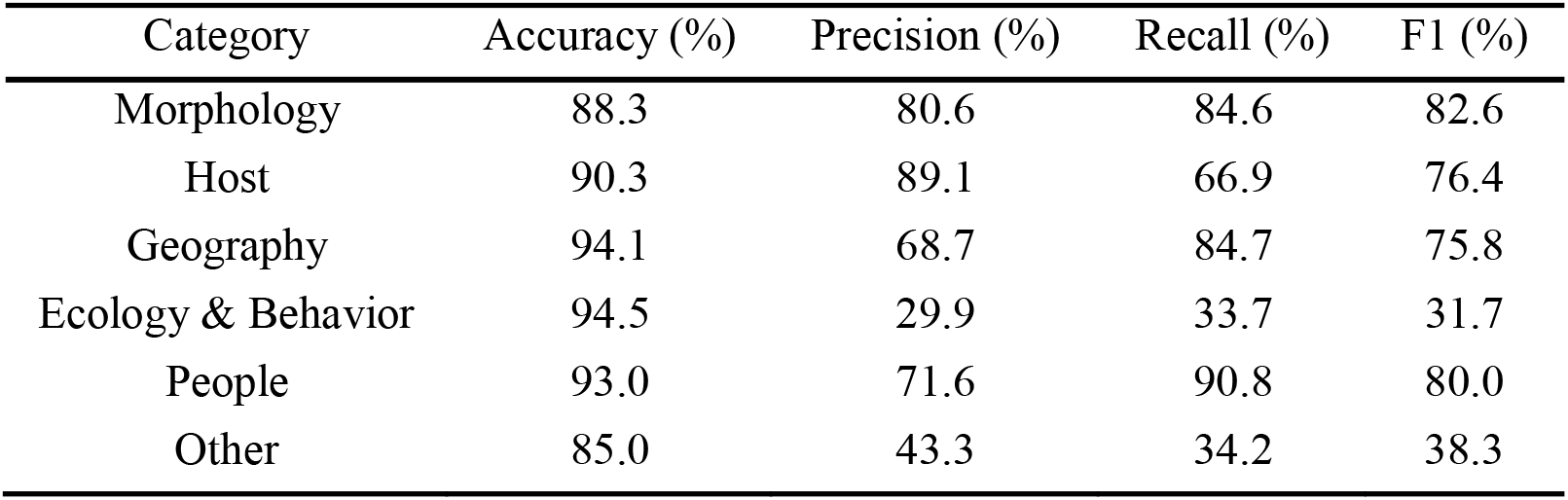
Accuracy and classification performance of Human-based and LLM-based labeling. Overall accuracy, average accuracy per species, and category-specific accuracy rates between Human-based and LLM-based labeling. Additionally, F1 scores, precision, and recall for each category are shown.

| Category | Accuracy (%) | Precision (%) | Recall (%) | F1 (%) |
| --- | --- | --- | --- | --- |
| Morphology | 88.3 | 80.6 | 84.6 | 82.6 |
| Host | 90.3 | 89.1 | 66.9 | 76.4 |
| Geography | 94.1 | 68.7 | 84.7 | 75.8 |
| Ecology & Behavior | 94.5 | 29.9 | 33.7 | 31.7 |
| People | 93.0 | 71.6 | 90.8 | 80.0 |
| Other | 85.0 | 43.3 | 34.2 | 38.3 |

The total count in each category was similar between Human-based and LLM-based labeling (Fig. 1). In Human-based labeling, the major categories were *Morphology* (894 species, 32.7%), *Host* (646 species, 23.6%), and *People* (534 species, 19.5%). Similarly, in LLM-based labeling, *Morphology* (940 species, 34.3%), *Host* (486 species, 17.7%), and *People* (534 species, 19.5%) were frequently classified.

**Fig. 1.**
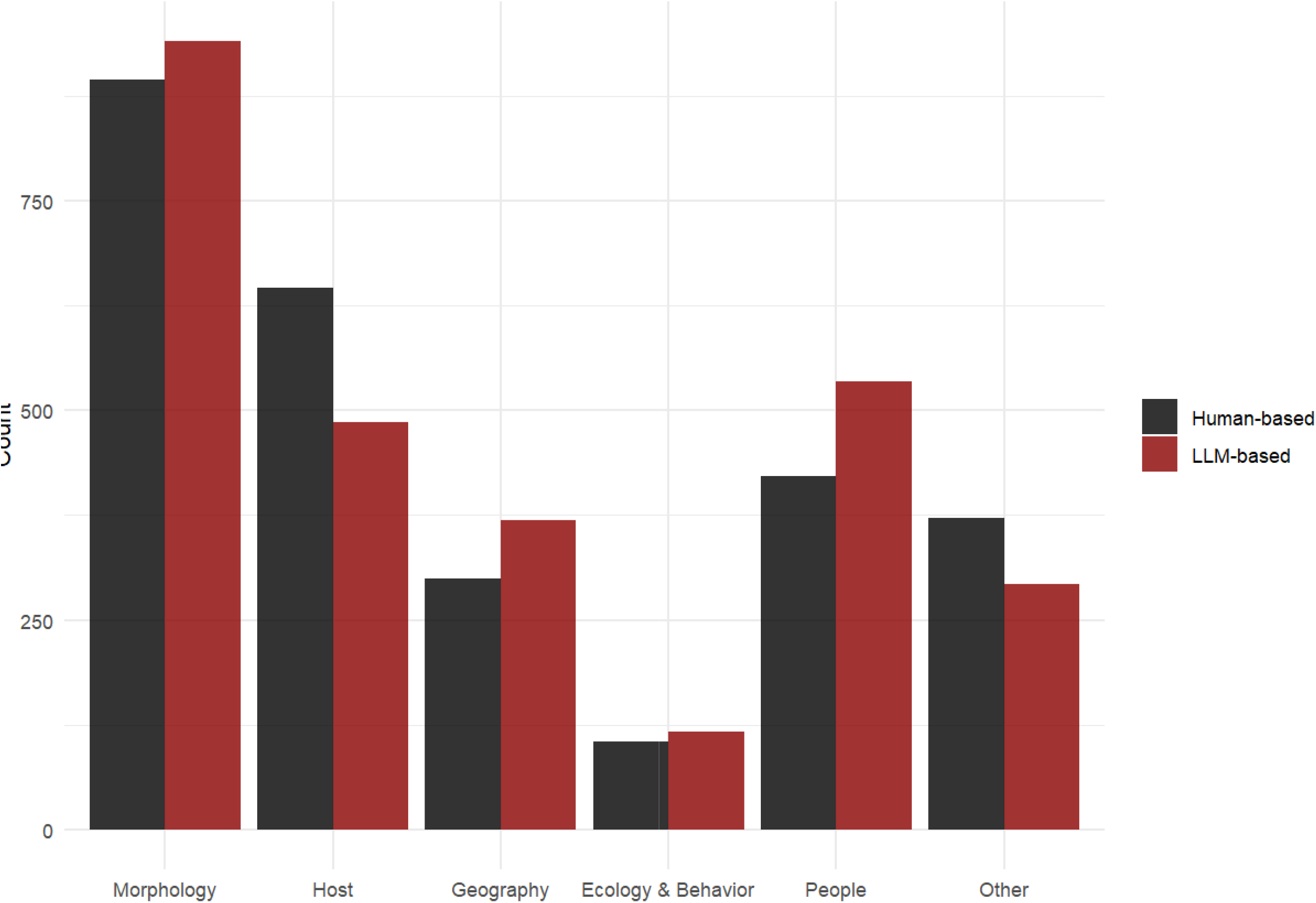
Total count of species classified into each category by Human-based and LLM-based labeling. Human-based labeling (black bars) and LLM-based labeling (dark red bars) classified nearly the same number of species in each category.

Similar classification tendencies in *Morphology, Host, Geography*, and *People* were observed (Fig. 2). High PCCs were found in *Morphology* (r = 0.951), *People* (r = 0.947), *Host* (r = 0.936), and *Geography* (r = 0.904). However, *Ecology & Behavior* (r = 0.431) and *Other* (r = 0.674) had lower PCCs. MAE values across all categories were small, remaining below 1. Human-based and LLM-based classifications showed similar regression curves in GAM analysis for category comparisons (Fig. 3). LLM-based labeling effectively detected the overall decreasing trends in *Morphology* and *Host*. In Human-based classification, *Morphology* exhibited a trend shift around 1900, but this was not reflected in the LLM-based smoothed curve. Additionally, differences in the rate of decline in *Host* were observed between the two methods.

**Fig. 2.**
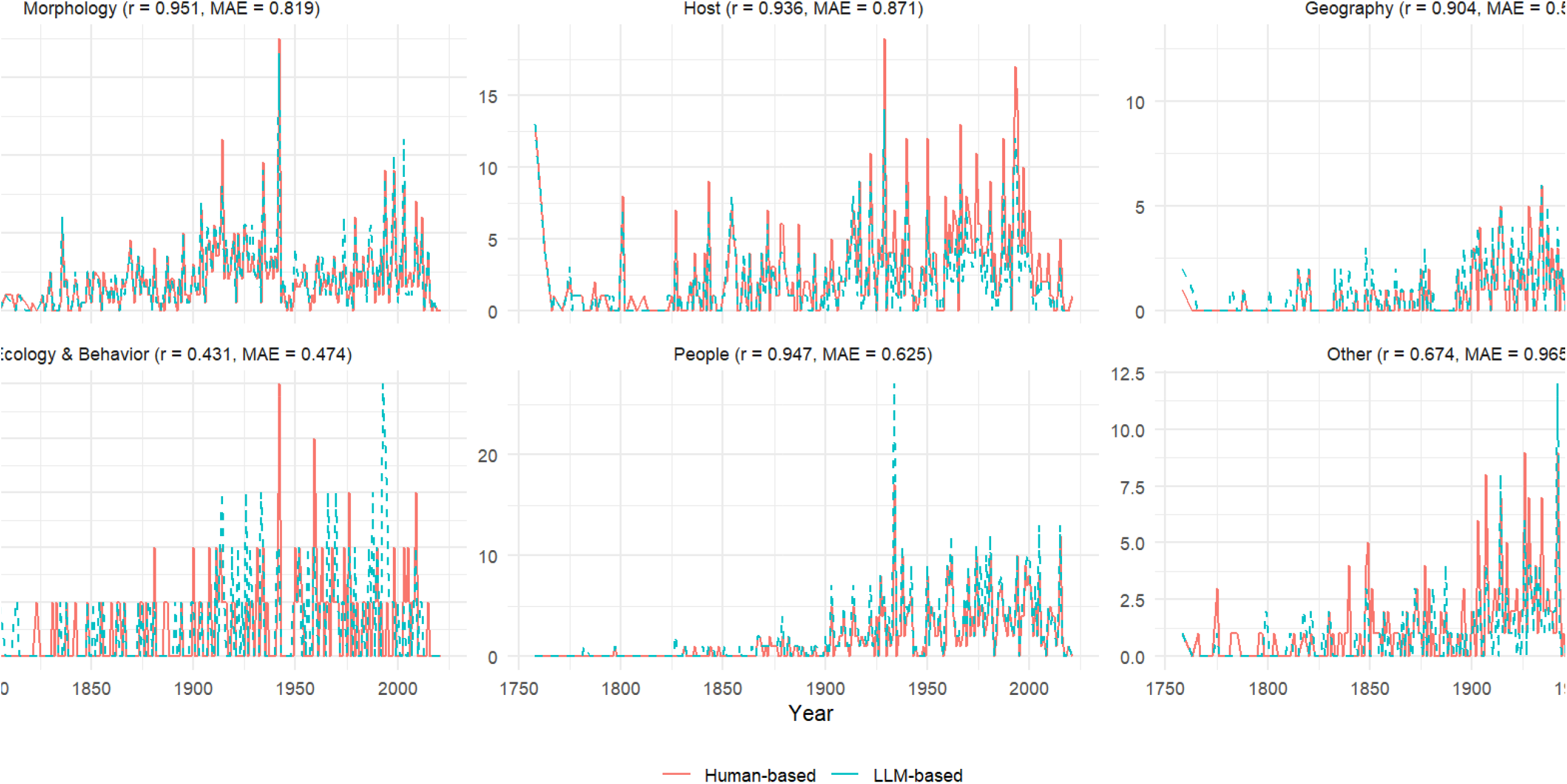
Temporal shifts in the total count of species classified into each category per year. Trends in species classification by Human-based labeling (solid dark red line) and LLM-based labeling (light blue dashed line) are shown. Both methods exhibited similar temporal trends.

**Fig. 3.**
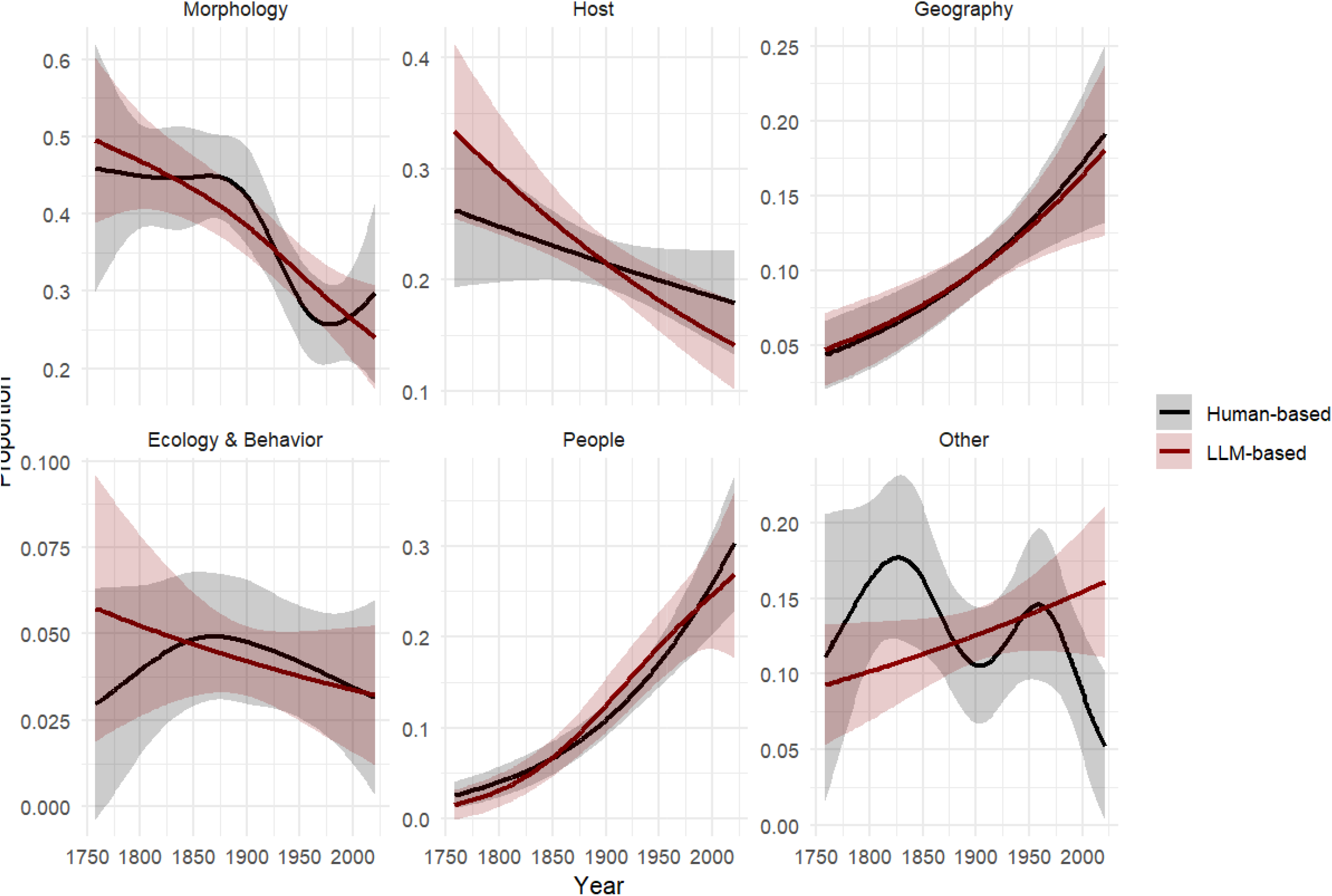
Smoothed regression curves with 95% confidence intervals in GAM for Human-based and LLM-based labeling. Human-based labeling (black) and LLM-based labeling (dark red) results are compared. *Geography* and *People* exhibited nearly identical curves, while *Morphology* and *Host* showed similar decreasing trends. In contrast, *Ecology & Behavior* and *Other* displayed different patterns.

Fig. 4 shows temporal changes in classification rates using GAM. The LLM-based smoothed curve showed a trend similar to that of the Human-based smoothed curve. Both classifications indicated a decreasing trend in *Morphology*, with a particularly sharp decline around 1900 in Human-based classification. Conversely, *Geography* and *People* exhibited an increasing trend in both methods.

**Fig. 4.**
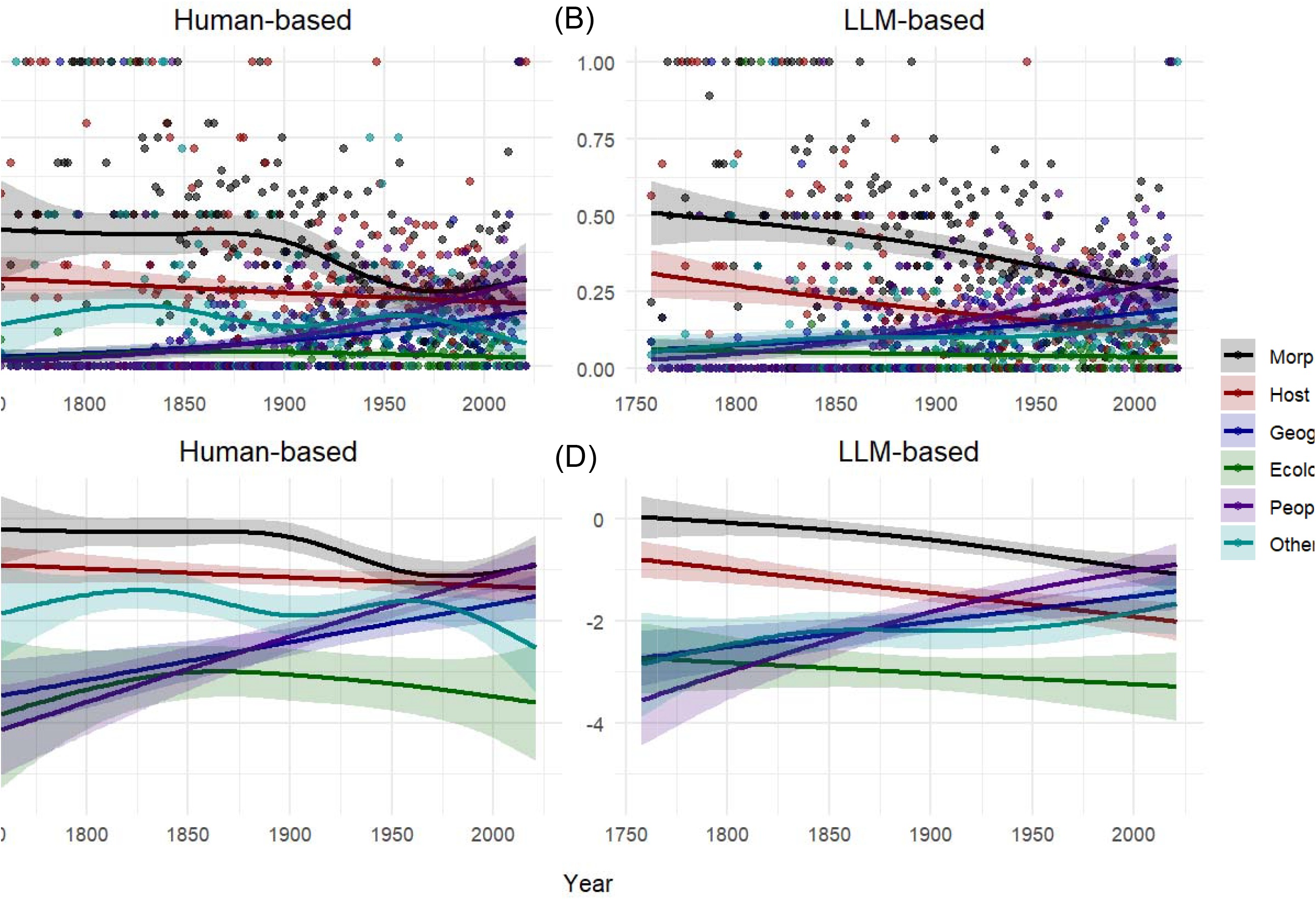
Temporal trends expressed by regression curves with 95% confidence intervals in GAM. Categories are color-coded as follows: *Morphology* (black), *Host* (dark red), *Geography* (dark blue), *Ecology & Behavior* (dark green), *People* (Purple), and *Other* (light blue). (A) Temporal trends in the proportional use of etymologies across different categories classified by Human-based method. Each dot represents an observation, and the smooth lines depict trends with 95% confidence intervals. (B) Smooth terms for temporal variations in GAM across different categories classified by Human-based methods. (C) Temporal trends in the proportional use of etymologies across different categories based on LLM-based labeling. (D) Smooth terms for temporal variations in GAM across different categories based on LLM-based labeling.

## DISCUSSION

In this study, the accuracy of LLM-based classification was evaluated using Human-based classification as ground-truth data. The analyses revealed that LLM-based labeling can reflect the general temporal trends in most categories of phytophagous arthropods. It demonstrated the potential of LLM to reduce the substantial labeling costs associated with traditional methods significantly.

In particular, *Morphology, Host, Geography*, and *People* achieved F1 scores above 75% and accuracy above 90%, indicating high classification performance. On the other hand, *Ecology & Behavior* and *Other* exhibited extremely low F1 scores, indicating instability in classification. These results are consistent with previous research(Inoshita et al., 2025), which also reported low accuracy in *Ecology & Behavior* and *Modern & Past Culture*. This may be due to the insufficient availability of data related to *Ecology & Behavior*, limiting the model’s ability to grasp deep contextual meaning.

PCCs for *Morphology, Host, Geography*, and *People* exceeded 0.9, showing strong correlations. This suggests that LLM-based classification closely resembles Human-based classification. In LLM-based labeling, *Morphology* and *Host* showed a decreasing trend, while trends, analyses using GAM, *Geography*, and *People* exhibited similar trends in both methods. These findings align with previous research on spiders, and rotifers(Macêdo et al., 2023; Mammola et al., 2023), suggesting that the decline of Morphology-related names and the rise of Geography and People-related names are general naming trends across taxa. Even though general trends resembling Human-based labeling were observed in LLM-based labeling, subtle differences were present in *Morphology*. Human-based classification indicated that *Morphology* followed different trends before and after 1900, a shift that LLM-based classification did not detect. In conclusion, LLM-based labeling is suitable for long-term trend analysis, but subtle changes may be overlooked.

This drastic decline in *Morphology* may be attributed to the rediscovery of Mendel’s work and the development of ultrastructural techniques (e.g., electron microscopy). These advancements in biology may have elevated taxonomic descriptions to higher levels, introducing genetic and ultrastructural perspectives into the taxonomy and reducing the emphasis on classical macromorphological naming. The increase in *Geography* and *People* may be interpreted as a relative effect of the decline in *Morphology*. However, there may be underlying factors influencing these trends. Therefore, more detailed investigations into the etymology of individual species are required.

Regarding eponyms, some species have been named after universally offensive people (e.g., *Hypopta mussolinii*, Turati 1927; *Rochlingia hitleri*, Guthörl 1934; *Anopthalmus hitleri*, Scheibel 1937). Additionally, 89.4% of the eponyms in Mollusks honored men [Vendetii], highlighting a gender imbalance in eponyms. Beyond these ethical concerns, some taxonomists argue that naming conventions rooted in Latin (though no longer required in ICZN) marginalize indigenous languages and cultures(Gillman and Wright, 2020; Wright and Gillman, 2022). Given these considerations, calls for ICZN revision have emerged (Gillman and Wright, 2020; Roksandic et al., 2023; Rummy and Rummy, 2021; Wright and Gillman, 2022). However, in terms of scientific consistency, opposing views on ICZN revision have also been voiced (Antonelli et al., 2023; Ceríaco et al., 2023; Jiménez-Mejías et al., 2024; Orr et al., 2023; Pethiyagoda, 2023; Thiele, 2023). The ongoing debates surrounding scientific nomenclature are highly complex and remain unresolved. This study identified an increase in eponyms, which should be taken into account in these discussions.

Furthermore, species named after their host are often selected as the subjects for HAD studies(Mlynarek et al., 2023). Thus, the observed decline in *Host* may suggest that host-association studies tend to focus on species named in the past, potentially introducing bias. In conclusion, scientific names may influence not only social and ethical issues but also the advancement of science itself.

The LLM classified the origins of epithets with high accuracy in just one hour. This result is consistent with previous research(Inoshita et al., 2025), confirming that LLM-based labeling is applicable to arthropods beyond spiders. Although *Ecology & Behavior* cannot yet be classified with high accuracy, time and effort can be saved by applying cross-validation specifically to this category. Further evaluation of LLM-based classification across other taxa is required. Moreover, this study provides a novel perspective on the social and scientific discussion surrounding scientific names by elucidating naming trends in phytophagous arthropods. Understanding naming trends in animals will lay the foundation for scientific discussions and contribute to the development of better nomenclature rules. The integration of LLM into taxonomy will enhance our understanding of species, their nature, and their history.

## ACKNOWLEDGMENTS

This study was conducted using the dataset published by Heard et al. (2023) in “Can species naming drive scientific attention? A perspective from plant-feeding arthropods”. We sincerely appreciate their efforts in collecting and sharing the data, which provided a valuable foundation for this study. We also thank Takumi Taga for his helpful comments on the manuscript.

## AUTHOR CONTRIBUTIONS

KN designed the study, performed data analyses, and wrote the manuscript. KI developed the prompt for the LLM and prepared the data for analysis. KI and HS made significant contributions to editing the drafts. All authors approved the final version of the manuscript.

